# New Space-Time Tradeoffs for Subset Rank and *k*-mer Lookup

**DOI:** 10.64898/2026.03.16.712042

**Authors:** Anastasia C. Diseth, Simon J. Puglisi

## Abstract

Given a sequence *S* of subsets of symbols drawn from an alphabet of size *σ*, a subset rank query srank(*i, c*) asks for the number of subsets before the *i*th subset that contain the symbol *c*. It was recently shown (Alanko et al., Proc. SIAM ACDA, 2023) that subset rank queries on the spectral Burrows-Wheeler lead to efficient *k*-mer lookup queries, an essential and widespread task in genomic sequence analysis. In this paper we design faster subset rank data structures that use small space—less than 3 bits per k-mer. Our experiments show that this translates to new Pareto optimal SBWT-based *k*-mer lookup structures at the low-memory end of the space-time spectrum.

## 1 Introduction

The problem of *k*-mer lookup, which is the task of quickly determining if a query *k*-mer is a member of a set of *k*-mers and returning the rank of the *k*-mer if found, is central to many modern genomic-analysis pipelines [16] and processes, such as pseudoalignment [4, 9, 12, 11]. Common solutions are based on hashing and minimizers [19, 17], or make use of the spectral Burrows-Wheeler transform (SBWT) of the *k*-mer set [3]. In this paper, we focus on a key data structural problem at the heart of SBWT-based k-mer lookup: efficient *subset rank queries*.

Given an alphabet Σ of *σ* symbols, and a sequence *S* of subsets of Σ, a *subset rank* query subset-rank(*i, c*) asks for the number of subsets before the *i*th subset in *S* that contain the symbol *c*. For example, if Σ = A, C, G, T and *S* = {T}{G}{A, C, G, T}{}{}{C, G}{}{A}{}{A}{A, C}{}{}{A}{A}, then we would have subset-rank_*S*_(8, A) = 2, subset-rank_*S*_(12, A) = 4, and subset-rank_*S*_(8, G) = 3.

Alanko et al. [3] recently showed that a series of at most *k* pairs of subset rank queries over the spectral Burrows-Wheeler transform of an input string (or set of input strings) *T* is enough to determine the colexicographic rank of a query *k*-mer in *T* ‘s *k*-mer spectrum, and thus perform *k*-mer lookup. The resulting SBWT-based *k*-mer lookup indexes are fast compared to widespread hash-based schemes [19, 17] and encode the *k*-mer spectrum efficiently.

Beyond *k*-mer lookup, subset rank queries are an efficient means by which to navigate the *k*-order de Bruijn graph corresponding to the *k*-mer spectrum encoded in the SBWT, and thus have a broad array of applications. Examples include, efficient computation of finimizers [6], longest common *k*-mer suffixes and *k*-bounded matching statistic [1, 6], the implementation of variable-order de Bruijn graphs [7, 8], and approximate *k*-mer lookup. Thought of another way, the SBWT efficiently encodes a truncated suffix tree [1] of *T*, and subset rank queries then implement the child operation at nodes, or continued decent along edges and are thus of immense interest.

In the original SBWT paper [3], Alanko et al. describe four subset rank schemes with different space-time tradeoffs. The smallest structure uses just 2.3 bits per *k*-mer in practice, but is at least an order of magnitude slower than their fastest method, which uses around 4.3 bits per *k*-mer.

In this paper we drastically flatten the trade-off curve for subset rank queries, describing small subset rank structures with query times approaching those of the bigger, faster methods. We achieve these gains through both improved data structure design and better engineering of internal components.

The remainder of this paper is structured as follows. In the next section we cover basic concepts and related work. Section 3 is the body of the paper, where we describe our new data structures for subset-rank, including a microbenchmark to prior subset-rank structures. Section 4 then reports on experiments applying our new structures to the *k*-mer lookup problem, comparing the original SBWT-based methods from Alanko [3].

## 2 Preliminaries

We use the usual array notation for strings throughout. We consider a string *T* of length |*T*| = *n* as a sequence of *n* symbols *T* [0..*n* − 1] = *T* [0]*T* [1] … *T* [*n* − 1]. The empty string is denoted *ϵ* and |*ϵ*| = 0.

In this paper we are particularly interested in strings on base-4 and base-2 (i.e. binary) alphabets. In examples, we sometimes use the DNA alphabet Σ = {A, C, G, T} and *σ* = |Σ| = 4. When describing algorithms and data structures, we assume a minimal base-4 alphabet Σ = {0, 1, 2, 3}.

The *substring* of *T* starting at symbol *i* and ending at symbol *j* is denoted *T* [*i*..*j*]. We also use the half-open interval notations *T* (*i*..*j*] = *T* [*i* + 1..*j*] and *T* [*i*..*j*) = *T* [*i*..*j* − 1]. A *prefix* is a substring starting at position 0 and a *suffix* is a substring ending at position *n* − 1. The *colexicographic order* of two strings corresponds to the lexicographic order of their reverse strings. We define a *k*-mer as a string of length *k*.

The set of distinct *k*-mers occurring in a string *T* is referred to as the *k-spectrum* of *T*.

### Definition 1

*(k-Spectrum)* The *k*-spectrum of a string *T*, denoted with *S*_*k*_(*T*), is the set of all distinct *k*-mers in the string *T*′ = $^*k*−1^*T*. The *k*-spectrum *S*_*k*_(*T*_1_, …, *T*_*m*_) of a set of *m* strings *T*_1_, …, *T*_*m*_ is the union 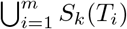.

The $s are for technical convenience to ensure there is a *k*-mer ending at every position in *T*.

By way of illustration, consider the string *T* = TAGCAAGCACAGCATACAGA, for which we have *S*_3_(*T*) = {AAG, ACA, AGA, AGC, ATA, CAA, CAC, CAG, CAT, GCA, TAC, TAG $$$, $$T, $TA}.

In this paper the primary operation we aim to support is the *k-mer lookup* query, defined as follows.

### Definition 2

*(k-mer lookup query)* Given an input a string *P* of length *k*, a *k*-mer lookup query returns the colexicographic rank of *P* in *S*_*k*_(*T*), or ⊥ if *P* ∉ *S*_*k*_(*T*)

The Spectral Burrows-Wheeler transform (SBWT) [3] transforms a *k*-spectrum into a sequence of subsets of the alphabet. Conceptually simple operations on the SBWT subset sequence allow *k*-mer lookup queries to be efficiently supported. Although it is not needed to understand our subset rank data structures, for completeness we now define the SBWT and give a brief overview of its salient properties, referring the reader to [3, 1, 5] for a more extensive treatment.

### Definition 3

*(Spectral Burrows-Wheeler Transform)* Let *S*_*k*_ be a *k*-spectrum and let 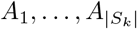 be the colexi-cographically sorted elements of *S*_*k*_. The SBWT is the sequence of sets of characters 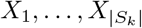 with *X*_*i*_ ⊆ Σ such that *X*_*i*_ = ∅ if *i >* 1 and *A*_*i*_[1..*k* − 1] = *A*_*i*−1_[1..*k* − 1], otherwise *X*_*i*_ = {*c* ∈ Σ | *A*_*i*_[1..*k* − 1]*c* ∈ *S*_*k*_}.

The *k*-mers from the 3-spectrum *S*_3_(*T*) above, listed in colexiographical order are: $$$, CAA, ACA, GCA, AGA, $TA, ATA, CAC, TAC, AGC, AAG, CAG, TAG, $$T, CAT; and the SBWT is the sequence of sets: {T}, {G}, {ACGT}, ∅,∅, {CG},∅, {A}, ∅, {A}, {AC}, ∅, ∅, {A }, {A }.

The SBWT set represent the labels of outgoing edges in the node-centric de Bruijn graph, such that only outgoing edges from *k*-mers that have a different suffix of length *k* − 1 than the preceeding *k*-mer in the colexicographically sorted list are included. This gives the SBWT sequence an important *balance property*, namely that |*S*_*k*_(*T*)| − 1 = ∑ |*X*_*i*_|. That is, the number of *k*-mers in the spectrum is exactly one greater than the total sizes of the sets. Moreover, on typical genomics use cases, singleton subsets dominate, consistuting around 95% of all subsets.

The SBWT encodes the set of *k*-mers *S*_*k*_(*T*) as a colexicographically sorted list. In the context of *k*-mer lookup, the SBWT allows us to navigate that ordered list via the operation *ExtendRight*, defined as follows.

### Definition 4

*(ExtendRight)* Let [*s, e*]_*α*_ be the *colexicographic interval* of string *α*, where *s* and *e* are respectively the colexicographic ranks of the smallest and largest *k*-mer in *S*_*k*_ suffixed by the substring *α. ExtendRight*([*s, e*]_*α*_, *c*) denotes the *right extension* of the interval [*s, e*]_*α*_ with a character *c* ∈ Σ. This outputs the interval [*s*′, *e*′]_*αc*_, or ⊥ if no such interval exists.

As explained in [3], *ExtendRight* can be answered via using two *subset rank* operations on the SBWT. We first define the *rank* operation. Let *B* be a bit string (or bit vector) of length *n*, for every index *i* ≤ *n* and *b* ∈ {0, 1}, rank_*b*_(*B, i*) is equal to the number of *b*-bits in *B*[0..*i*).

### Definition 5

*(Subset rank)* Let *X*_1_, …, *X*_*n*_ be a sequence of subsets of symbols from alphabet Σ. A subset rank query subset-rank(*i, c*) for index *i* and symbol *c* ∈ Σ returns the number of subsets *X*_*j*_ with *j < i* such that *c* ∈ *X*_*j*_.

With this, ExtendRight([*s, e*]_*α*_, *c*) can be computed using the formulas *s*′ = 1 + *C*[*c*] + subset-rank(*s* − 1, *c*) + 1 and *e*′ = 1 + *C*[*c*] + subset-rank(*e, c*), where *C*[*c*] is the number of occurrences of characters smaller than *c* in the SBWT sequence [3]. In our example *C*[A] = 0, *C*[C] = 6, *C*[G] = 9 and *C*[T] = 12.

## 3 Prior Work on Subset Rank Queries

The subset sequence of the SBWT can be represented in many ways to support subset rank queries. In this section, we provide an overview of three different data structures from [3], which we also use as performance baselines in our experiments. Our treatment here is not intended to be exhaustive, but rather to highlight that an interesting range of space-time tradeoffs are possible for subset rank queries. We use the notation from the previous section, that is, we are indexing a sequence *X*_1_, … *X*_*n*_ of subsets of a base-4 alphabet Σ = {0, 1, 2, 3}. A subset rank query takes as input an index *i* and a character *c* ∈ Σ, and returns the number of subsets *X*_*j*_ with *j < i* such that *c* ∈ *X*_*j*_.

### 3.1 Matrix Representation

This data structure uses a binary matrix *M* of size 4*n*, such that *M* [*i*][*j*] = 1 iff subset *X*_*j*_ contains the *i*-th symbol of the alphabet. The rows of the matrix are indexed for fast succinct rank queries. The subset rank query subsetrank(*j, c*) is answered with a binary rank query rank(*M* [*c*], *j*) on row *M* [*c*] up to index *j*. See Figure 1.

**Figure 1.**
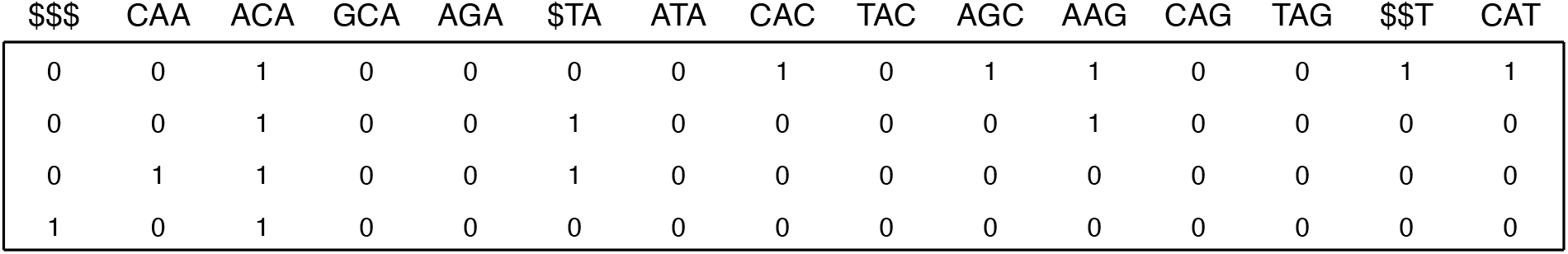
The Matrix representation of the SBWT for TAGCAAGCACAGCATACAGA with *k* = 3.

According to experiments in [3], this data structure is currently the fastest in practice. Their implementation uses the binary rank structure from [10]). The space used is ~4.3 bits per *k*-mer: 4 bits for the bits of the matrix column, plus some overhead for the binary rank structures on the rows.

### 3.2 Split Representation

The next—so-called *Split*—representation aims to exploit the property, stated in Section 2, that many of the SBWT subsets are singleton —in genomics use cases, typically 95% of SBWT subsets are singletons [3, 2].

Let *M*^−^ be the submatrix of matrix *M* in the matrix representation that contains only the columns of *M* with exactly one 1-bit set, and let *M* ^+^ be the submatrix of *M* containing the rest of the columns. Let *B* be a bit vector of length equal to the number of columns in *M*, marking with 1-bits which columns of *M* are in *M* ^+^.

We index both *M* ^+^ and *M*^−^ for character rank queries. Matrix *M* ^+^ is indexed like in the matrix representation. Matrix *M*^−^ is replaced by a string *W* that is the concatenation of the labels corresponding to the bits in the columns. The string *W* is indexed for base-4 rank queries (in [3], a wavelet tree is used). *B* is indexed for rank queries. Because singletons are expected to dominate, *B* will be sparse and is efficiently indexed using Elias-Fano encoding (see, e.g., [18, 15]). The final index consists of *B, M* ^+^ and *W* and rank support structures. See Figure 2.

**Figure 2.**
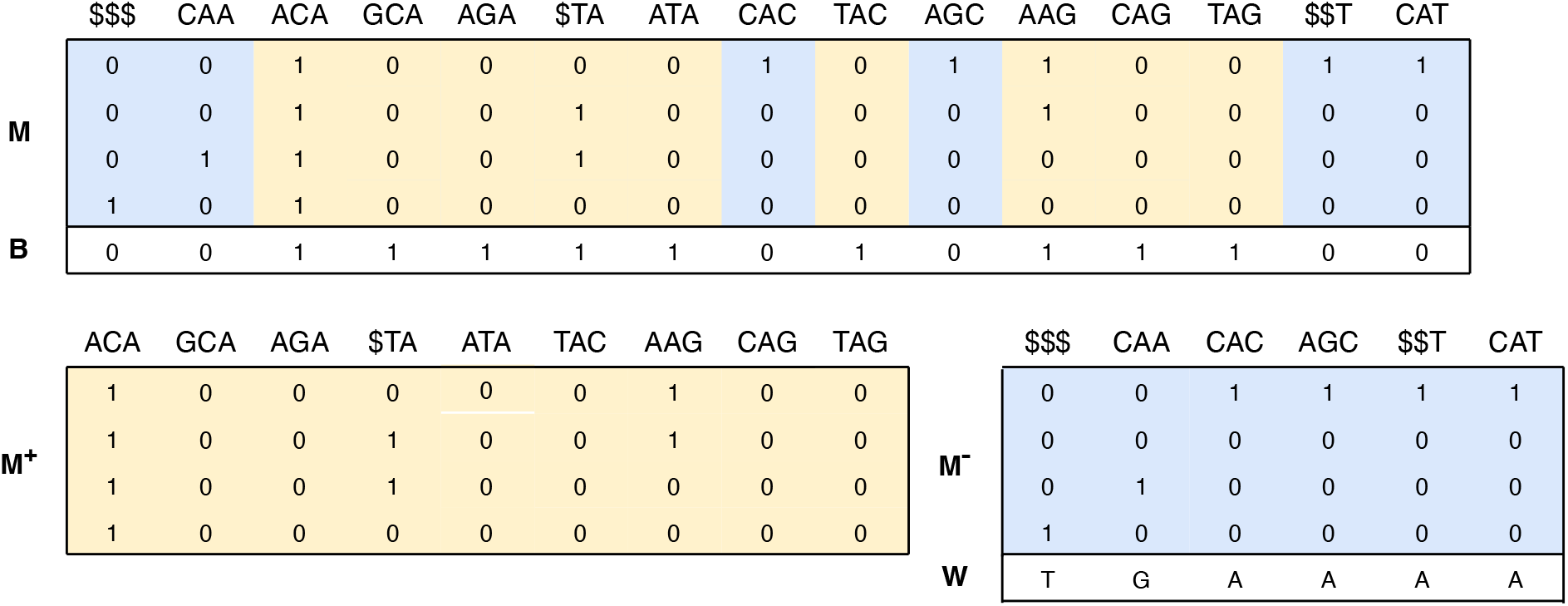
The Split representation of the SBWT for TAGCAAGCACAGCATACAGA with *k* = 3.

A query subset-rank(*i, c*) can now be answered by using a rank query on *B* to determine how many of the first *i* columns of *M* went to *M* ^+^ and *M*^−^, then using the rank structures of *M* ^+^ and *M*^−^ to count the number of characters in the corresponding prefixes of columns in both matrices, and returning the sum of the two counts. That is, if *r* = *rank*_1_(*B, i*), then answer to the query is *rank*_1_(*M* ^+^[*c*], *r*) + *rank*_*c*_(*W, i* − *r*).

Split uses around ~2.6 bits per *k*-mer and is 3-4 times slower than matrix.

### 3.3 Concat Representation

This data structure, dubbed Concat, uses a concatenation of the contents of the subsets, and an encoding of the sequence of sizes of the subsets. In more detail, let *S*(*X*_*i*_) be the concatenation of the characters in subset *X*_*i*_. If *X*_*i*_ is the empty set, we define *S*(*X*_*i*_) = $, i.e., a special symbol not in the base-4 alphabet. We build the string *L* = *S*(*X*_1_)*S*(*X*_2_) … *S*(*X*_*n*_) and index it for rank queries. The empty sets are represented as $s to be able to encode the sizes of the subsets with a bit vector *B* that is the concatenation 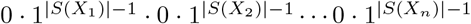. If the sequence of subsets contains a large number of singleton or empty sets, the bit vector *B* is sparse, and is efficiently stored using Elias-Fano encoding (see, e.g., [18, 15]). The bit vector *B* is indexed for select-0 queries.

The query subset-rank(*i, c*) is then answered by computing rank_*c*_(*L*, select_0_(*B, i* + 1) − 1). See Fig. 3. Concat uses around ~2.3 bits per *k*-mer and is 40-50 times slower than matrix.

**Figure 3.**
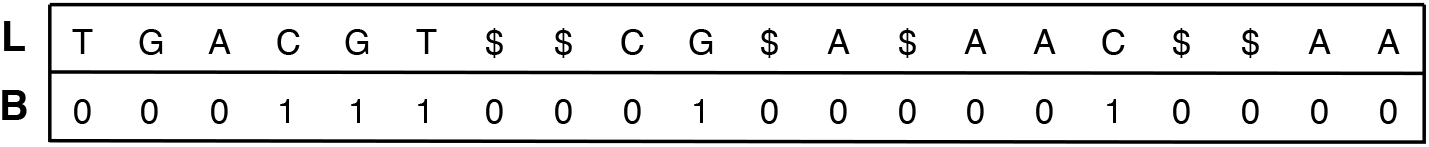
The Concat representation of the SBWT for TAGCAAGCACAGCATACAGA with *k* = 3.

## 4 Better Building Blocks

The efficiency of the Split and Concat schemes from [3] described in the previous section depend critically on the efficiency of the structures used for 1) rank over sparse integer sets and 2) rank over base-4 sequences. For the former, the implementations in [3] use an Elias-Fano (EF) implementation from Ma et al. [15]. For the latter, a wavelet tree (WT) as implemented in the SDSL lite [10] is used.

In this section we show that there are better choices for these internal data structures. We first describe an Elias-Fano-like structure in which the bucket size is fixed to 256, allowing various improvements in query performance, at the cost of space-optimality. We call this data structure Pred8. We then describe two fast alternatives to the WT for base-4 rank. The first is derived from Alanko et al. [2], where it is used to implement subset wavelet trees. A second, faster, method involves reordering the bits of a base-4 sequence, which has been contemporaneously described by Koerkamp in [13].

We show, via microbenchmarks, these structures lead to immediate improvements in query time for both the Split and Concat subset rank schemes, at a minor space cost, and may also be of independent interest.

### 4.1 Rank on Sparse Sets

We are given a set *S* of *m* integers *i*_0_ *< i*_1_ *<* … *< i*_*m*−1_ over a universe *U*, |*U*| = *u*. We will now describe a data structure—essentially a version of Elias-Fano (see, e.g., [15])— supporting rank and predecessor queries on *S*.

Let *L* be an array of *m* bytes where *L*[*x*] is the lower 8 bits of *i*_*x*_ ∈ *S*. We conceptually divide the universe into buckets of size 256, where the *j*th bucket, denoted *b*_*j*_, corresponds to the interval [256*j*..256(*j* + 1)). We store a bitvector *B* such at *B*[*j*] = 1 if and only if there is at least one element from *S* falling in bucket *b*_*j*_. We also store an array *H* of *u/*256 integers, one for each bucket, which contains indexes into *L*, defined as follows. Assume one or more elements from *S* fall into bucket *b*_*j*_ (i.e. *B*[*j*] = 1)and let *i*_*x*_ ∈ *S* be the smallest of those elements. We set *H*[*j*] = *x* indicating that *L*[*x*] is the first entry in *L* containing lower bits corresponding to elements of *S* falling in bucket *b*_*j*_. If, on the other hand, *b*_*j*_ is empty, then we set *H*_*j*_ = *y*, such that *i*_*y*_ is the predecessor of 256*j* in *S*; in other words, *i*_*y*_ is the largest element in *S* before the start of bucket *b*_*j*_.

Given a predecessor pred(*p*) query, we can determine the *x* such that *i*_*x*_ ∈ *S* is the predecessor of *p* with the above data structure as follows. If the bucket containing *p*, i.e., *b*_*j*_, *j* = *p/*256, is empty (determined by checking *B*[*j*]) then the answer to pred(*p*) is given by *H*[*j*]. Otherwise, *p*’s bucket is not empty. We calculate the number of elements in it as *c* = *H*[*j* + 1] − *H*[*j*] + *B*[*j* + 1] and we scan *L*[*H*[*j*]..*H*[*j*] + *c*] until we determine the predecessor of *p*, or, if *p* is smaller than all elements in the bucket we return *H*[*j*] − 1 as the index of the predecessor. Answering a rank query rank(*p*) is immediate given the index of the predecessor after checking if the predecessor is equal to *p*. When implementing the above scheme we use an array of bytes for *L* and array of 32-bit integers for *H*. We use the most significant bit of *H*[*i*] to store the bit *B*[*i*], thus limiting *m <* 2^31^. Having all elements aligned to byte and word boundaries, avoids the bit-picking present in usual EF implementations. Queries are very simple, involving just a single access to *H* followed by a possible scan of at most 256 bytes. The space used is *u/*8 + 8*m* bits. This compares with (*u/*256) + *m* + 8*m* bits for a traditional EF structure with bucket size set to 256, and *m*(2 + log ⌈*u/m*⌉) bits for an EF structure with the bucket size set for minimal space usage.

For sufficiently dense sets this structure gives an interesting practical space-time trade-off. For example, when the rate of 1 bits if 1*/*16, we have *m* = *u/*16 and our data structure takes 140*u/*256 bits, which, given the ease of querying, compares favorably with the 145*u/*256 bits taken by a traditional EF structure with bucket size 256 takes, and the 96*u/*256 bits of a minimal EF structure takes (ignoring space needed for select support in the latter two structures). This is illustrated in the top section of Table 1 below, where we compare versions of the Split data structure, one using Pred8 for rank on the set of non-singleton-set positions, and another using the EF data structure [15] as used in [3]. Using Pred8 is noticeably faster on all three data sets, without increasing space-usage.

**Table 1:** Time (*µ*secs/query) and space (bits per *k*-mer) for 1 million subset rank queries on data sets from Section 6.

|  | Ecoli |  | Salmonella |  | Human |  |
| --- | --- | --- | --- | --- | --- | --- |
|  | time | space | time | space | time | space |
| Split EF + wavelet tree | 0.36 | 2.67 | 0.46 | 3.73 | 0.48 | 3.51 |
| Split Pred8 + wavelet tree | 0.32 | 2.65 | 0.40 | 3.72 | 0.41 | 3.51 |
| Split Pred8 + natural packing | 0.26 | 2.77 | 0.40 | 3.84 | 0.41 | 3.63 |
| Split Pred8 + hi-lo packing | 0.25 | 2.77 | 0.34 | 3.84 | 0.35 | 3.63 |

### 4.2 Base-4 Rank

As already mentioned, the implementation of Split in [3] makes use of a wavelet tree for supporting rank queries over the base-4 sequence *W*. A balanced wavelet tree on *W* takes 2*n* bits and reduces a rank query on *W* to two rank queries on bit vectors. Unfortunately, however, those two rank queries are, in general, non-local, requiring access to two parts of the data structure that are distant in memory.

In [2], Alanko, et al. describe the following data structure for base-4 rank that performs rank on the base-4 input sequence—*W* in our case—more directly with the aim of reducing non-local memory accesses. *W* is divided into *n/b* blocks of *b* contiguous symbols. Let *W*_*i*_ denote the *i*th block, corresponding to symbols *W* [*ib*..(*i* + 1)*b*). We precompute, for each block, four *boundary ranks R*_*i*_[0..3], one for each possible base-4 symbol, such that *R*_*i*_[*c*] is the number of occurrences of symbol *c* in *W* [0..*ib*). 32-bits are used for each element in *R*_*i*_ so that *R*_*i*_ takes 2 words in total. The representation for block *W*_*i*_ is *R*_*i*_ followed by *W* [*ib*..(*i* + 1)*b*) packed into 2*b* bits. We describe two possible packings of symbols in Section 4.2.1 below, the first is from [2] and the second is new.

To compute rank_*c*_(*W, i*) we access the *i/b*th block representation and scan the symbols it contains, counting occurrences of symbol *c* up to position *i* mod *b*. We then add this count to *R*_*i/b*_[*c*] to complete the answer. Block representations are of fixed size, so accessing one takes constant time if we store them consecutively in memory. The overhead incurred by the boundary ranks is 4 · 32 = 128 bits per block, or 128*/b* bits per symbol. Setting *b* = 2^9^, which is the block size in the experiments presented in this paper, the overhead incurred by the boundary ranks is 4 · 32*/*2^9^ = 0.25 bits per symbol, and the structure takes 2.25 bits per symbol in total.

#### 4.2.1 Symbol Packings

The natural way to represent symbols within a block in 2 bits per symbol is to pack 32 consecutive symbols into a 64-bit word, so that, for example, the base-4 sequence

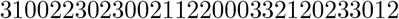

consisting of 32 symbols, would be represented as the 64-bit word

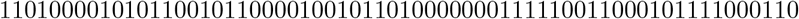

To compute the number of occurrences of symbol 2 inside a word, we first perform a bitwise & operation on it with the 64-bit binary mask

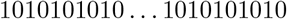

The count of 1 bits in the masked word gives the count of symbols greater than or equal to 2. Shifting the masked word one position to the right and using that as a new mask for the original word will leave a 1 bit set for each occurrence of symbol 3. The number of occurrences of symbol 2 inside the word is given by the count of occurrences of symbols greater than or equal to 2 subtracted by the number of occurrences of symbol 3. The computation of number of symbols smaller than 2 is identical, only the word is first negated.

A variation on the above scheme is to store the bits in a word slightly differently, so that all the high bits of the symbols come first, followed by all the low bits. In this way, the word above would be represented thus:

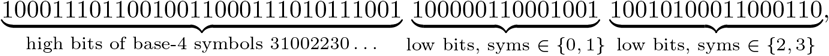

where the first 32 bits are the high bits of every symbol in the base-4 string, and the second 32 bits contain the low-bits of first the symbols having high bit equal to 0, followed by the low bits of those having 1 as the high-bit^1^.

This representation has the advantage that only bit shifts and popcount instructions are required to compute the in-word rank of any symbol—unlike in the first variant, where masking is required. In particular, to count the occurrences of a given symbol in the word, a popcount is performed on the 32 bit word containing the higher bits of the sequence. This count will determine how much the 32 bit word containing the lower bits needs to be shifted to the left or to the right depending on the symbol we are counting. A popcount on the correctly shifted lower bit word will give the number of occurrences of the symbol we are looking for.

For example, to count the number of occurrences of symbol 2 in the above sequence of 32 symbols, we first perform a popcount on the first 32-bit word containing higher bits.

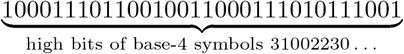

The number of 1-bits is 17. The second 32-bit word containing the lower bits is shifted 32 − 17 = 15 bits to the left.

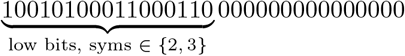

A popcount on the above word gives the number of occurrences of symbol 3, which is 7. The number of occurrences of symbol 2 is thus 17 − 7 = 10.

The lower part of Table 1 shows that Split is immediately improved by swapping in these approaches in place of wavelet trees. When Pred8 from the previous subsection is also used, the gain is more than 25% on all data sets. In the next section we show that much greater improvements are possible through completely new designs.

## 5 New Subset Rank Data Structures

In this section we present several novel data structures for subset rank support designed with SBWT and genomic scenarios in mind. Our overarching aims are to simultaneously improve memory locality and keep space usage low.

### 5.1 orrection sets

One drawback with the Split approach generally is that when answering a single subset rank query there are, essentially, three large and completely separate regions of memory that get accessed^2^: the top-level set that indicates single and non-singleton sets, the sequence of letters corresponding to singleton sets, and the non-singleton set representations. Each one of these three regions of memory is accessed essentially randomly, with three cache-misses likely incurred. Our approach in this section reduces the number of active memory regions to two per subset rank query, with the aim of reducing cache misses.

The data structure uses a concatenation of subsets, where each subset is represented by a single nucleotide letter. In addition, for each letter, we store a corresponding *correction set*, which is used to appropriately adjust a regular rank query on the concatenated string. In particular, let *U* (*X*_*i*_) be the lexicographically smallest character in subset *X*_*i*_, or the lexicographically smallest character in Σ if *X*_*i*_ = ∅. We build string *L* = *U* (*X*_1_)*U* (*X*_2_) … *U* (*X*_*n*_) of length *n*. The correction set for the lexicographically smallest character is the set of indices of the empty sets in *X*. For all other characters *c* in the alphabet (i.e. those not lexicographically smallest), we construct a correction set *Corr*_*c*_ containing the indices of subsets that contain *c* but were represented by a character other than *c* in *L*.

A subset rank query for character *c* at position *j* is answered by a base-4 rank query on *L* and a rank query on correction set *Corr*_*c*_ to find how many sets up to position *j* were not correctly represented in *L*. In other words, if *r* = rank_*c*_(*L, j*) and *x* = rank_1_(*Corr*_*c*_, *j*), then if *c* is the lexicographically smallest character in Σ, *x* is subtracted from *r*, otherwise the answer to be returned is the sum of *r* and *x*.

We now analyze the space complexity of the correction sets. Let *M* ^+^ be the set of subsets that are not of size 1, and let *m* = |*M* ^+^|. Due to the balance property of the SBWT (see Section 2)there must be a total of *m* elements in *M* ^+^. Let X_0_ be the set of empty sets in *M* ^+^, X_2_ the set of subsets containing 2 elements, X_3_ the set of subsets containing 3 elements, and so on, with X_*σ*_ the set of subsets containing *σ* elements. We obtain 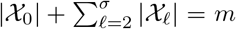 It is easy to see that for each set in X_2_, there must exist exactly 1 empty set, for each set in X_3_ exactly 2 empty sets, and so on. Thus 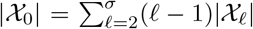. The correction set of the lexicographically smallest character is X_0_. Finally, each set *X*_*i*_ ∈ *M* ^+^, of length *ℓ* ≥ 2, must be included in *ℓ* − 1 correction sets, giving in total 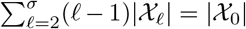. This gives a total of 2|X_0_| indices to be included in the correction sets. And since 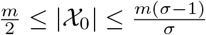 an upper bound for the total size of the correction sets is 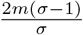.

In our implementation of this scheme, the correction set of nucleotide A is encoded using Pred8 data structure with block size 2^8^. The correction sets of nucleotides C, G, and T are much sparser and thus encoded using a version of Pred8 data structure that has block size 2^16^. See Figure 4.

**Figure 4.**
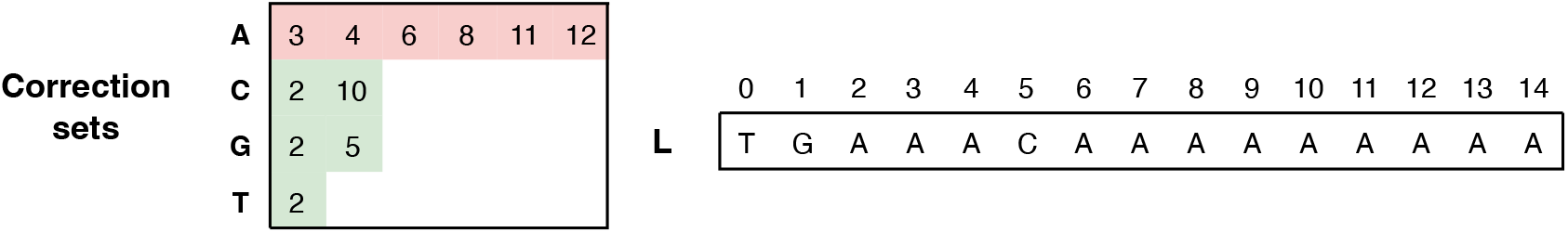
Correction sets scheme for SBWT: {T}, {G}, {ACGT}, ∅, ∅, {CG}, ∅, {A}, ∅, {A}, {AC}, ∅, ∅, {A}, {A}. For a subset-rank query for symbol A at position 12, there are 9 occurrences of A in *L* before position 12, and 5 positions in correction set of A less than 12, making the result 9 − 5 = 4.

An appealing property of the correction set approach is that the two queries involved in answering subset-rank(*i, c*)— on *L* and on *Corr*_*c*_—are independent of each other, allowing them to be performed in parallel if desired. This is in contrast to, e.g., Split, where the second and third parts of the query depend entirely on the first.

### 5.2 Blocked subset rank structures

Our next design attempts to reduce cache misses further. We divide the SBWT sequence *X* into *n/b* contiguous blocks containing *b* subsets each. For each block, we store *σ* counts, where count *C*_*j*_[*c*] is the number of subsets in *X* before subset *X*_*bj*_ containing letter *c*. In order to answer query subset-rank(*i, c*), we access block *j* = ⌊*i/b*⌋ and count the number of subsets before subset *i* (mod *b*) that contain letter *c*. We call this the *in-block rank*. We then add the in-block rank to *C*_*j*_[*c*] to complete the query.

Blocks are stored encoded. We denote with *E*_*j*_ the encoding for the *j*th block, which is comprised of sub-sets *X*_*jb*_, *X*_*jb*+1_, …, *X*_*jb*+*b*−1_. Each block is encoded separately and the encodings concatenated in memory *E* = *E*_0_*E*_1_*E*_2_ … *E*_*n/b*_. In general, the block encodings may be of non-uniform size, so in order to quickly locate the encoding of the *j*th block we store an array *P* of *n/b* block pointers, such that *P* [*j*] gives the starting position of block encoding *E*_*j*_ in the concatenation. The aim is that by choosing *b* appropriately, there is room in cache to hold both *P* and any one block encoding. Because *P* is relatively small and is accessed with every query, most of it should usually reside in cache and so the only cache misses likely incurred when answering subset-rank(*i, c*) come from accessing block encoding *E*_*i/b*_.

The precise encoding of the subsets comprising each block leads to different space-time tradeoffs. We now describe a simple encoding scheme. The block encoding *E*_*j*_ begins with the *C*_*j*_[*c*] counts (*σ* of them) defined above. These counts are stored as 32-bit integers. After these counts we store, using log *b* bits, the number of sets in the block that are not singletons (i.e. not of size 1). Then, each set that is not a singleton is encoded as follows. The block relative position is stored using log *b* bits and the set itself encoded using 4 bits, one bit per symbol marking if it is present in the set or not. The encoding of the non-singleton sets is followed by the encoding of the singleton sets packed as a concatenation of the symbols using 2 bits per symbol as in Section 4.2.1.

A subset-rank(*i, c*) query is answered as follows. Block *E*_*i/b*_ is accessed using the pointer at *P* [*i/b*] and the appropriate pre-block rank is read. Let *m*′ be the number of non-singleton positions in block *E*_*i/b*_ smaller than *i* mod *b*. We scan the non-singletons of the block sequentially to determine *m*′. During this scan, we inspect each subset encoding and keep count of the number of occurrences of symbol *c*. Having computed *m*′, we compute the rank of *c* in the packed symbol representation of the singleton sets at position *i* mod *b* − *m*′. The final answer to the rank query is the sum of the pre-block rank, non-singleton set rank, and singleton set rank.

We implemented several versions of the above scheme that differ principally in the way in which the non-singleton positions are encoded. In this paper we present two versions of Split with block sizes 2^8^ and 2^9^, each with its own space/time trade-off. In both versions non-singleton positions are encoded as differences between consecutive positions using 4 bits per position, with special treatment of larger gaps that would need more than 4 bits. As the number of gaps larger than 15 is bounded, using 4 bits per position instead of 8 resulted in more compact encodings. We also present a blocked version of the Correction Sets approach, where non-singleton sets in a block are encoded as positions of the correction sets for the given block. Encoding *E*_*i/b*_ stores, after the pre-block ranks, the sizes of the 4 correction sets within the block, followed by the correction sets in lexicographical order. Then follows a concatenation of the block’s SBWT sets (in one symbol per set) using the encoding of Section 5.1.

### 5.3 Fixed-block subset rank structures

With the aim of reducing average cache-misses per query even further, we now fix the block *encoding* size in the Correction Sets approach from Section 5.2. The idea is that each encoded block uses *e* words of space. *e* is chosen to incorporate most block encodings, with smaller encodings rounded up. If a block needs more than *e* words for its encoding, then the block encoding indicates this, and also where in memory to find the overflowing words. Encoding blocks this way eliminates the need for the block encoding pointers of the *P* array in the previous section: encoding *E*_*j*_ always starts at word *j* · *e*.

To answer subset-rank(*i, c*) we access block *i/b* directly at word *e* · (*i/b*). If the total encoding size of the block is less than *e* then all computation involves local information, otherwise we follow the stored pointer to obtain the other correction sets as required. Varying the overflow threshold *e* and the number of subsets per block *b* gives different space/time tradeoffs. A larger *e* allows for more blocks to be contained locally, but increases space usage as the block encoding is bigger. With smaller *b* space usage increases, as the average block overhead per SBWT subset becomes larger, but requires a smaller *e* to make a larger percentage of blocks below the overflow threshold.

## 6 Results

In this section, we report on experiments measuring the performance of our new subset rank data structures in comparison to existing data structures by Alanko et al [3]. Because our focus in this paper is on subset rank queries, and because subset rank has broader applications than *k*-mer lookup alone, we limit our experimental comparison to include only SBWT-based methods, and refer the reader to e.g., [3, 5] for a comparison of SBWT-based *k*-mer lookup to hash-based ones, such as [19], which may also be possible to further optimize.

### Methods

We measured the performance of the subset rank methods listed below.

- Plain Matrix and EF Matrix are implementations of the Matrix data structure from [3] described in Section 3.1 using, respectively, rankv5 [10] and modified Elias-Fano [15] for bitvector rank queries.
- Plain Split and EF Split are the Split method from [3] described in Section 3.1, with bitvector types as above. Pred8 Split is the fastest method from Section 4. The methods Blocked Split X are from Section 5.2 and represent blocks of size 2^*X*^ with the Split method.
- Correction Sets is the method of Section 5.1. Blocked Correction Sets is an implementation of the blocked approach from Section 5.2 using a block size of 2^8^. FB Correction Sets are three fixed-block implementations based on Section 5.3 with different *b* and *d* parameters. A and B have block size of 2^7^ and *e* = 6 and *e* = 7, respectively, while C has block size 2^6^ and *e* = 4.

### Data

We used three large genomic data sets with varying characteristics in our experiments.

- E.coli. 3,682 *E. coli* genomes having 249,991,499 distinct *k*-mers and 8,979,844 (3.6%) non-singleton sets^3^.
- Salmonella. 10,000 *S. enterica* genomes having 645,202,711 *k*-mers and 29,099,967 (4.5%) non-singleton sets^4^.
- Human. 60 *H. sapiens* genomes having 6,078,777,156 *k*-mers and 155,321,500 (2.6%) non-singleton sets^5^.

### Environment

Our experimental machine was a Dell PowerEdge C6420 with an Intel Xeon Gold 6248 CPU 2.50GHz CPU with 27.5 MB L3 cache shared by all cores, a 1MB L2 cache private to each core, and 64KB L1 data and instruction caches. The machine has 20 cores, but only a single thread of execution was used. The machine has 377GB of DDR RAM. The OS was AlmaLinux 8.10 and the compiler was g++ version 13.3 with the -O3 flag.

### Results and Discussion

We ran three different benchmarks, which we now describe and report on.

Single subset rank. This benchmark measures time taken by subset rank data structures for 1 million random subset rank queries. Query generation is not timed. The same set of queries is issued to each data structure.

Results are shown in the top half of Fig. 5. Considering methods using less memory than Plain Matrix, when memory is equated memory, the new methods are always faster than those from Alanko et al. [3]. That is, all previous small-space methods are dominated by the ones we describe, usually by a factor of 2 or more. Secondly, there is now a much smoother space-time tradeoff as space is increased, with the new methods approaching the speed of matrix as the amount of space they are allowed increases. Thirdly, blocked methods are consistently faster than their non-blocked counterparts. Finally, Corrections sets appears to have a slight advantage over Split.

**Figure 5.**
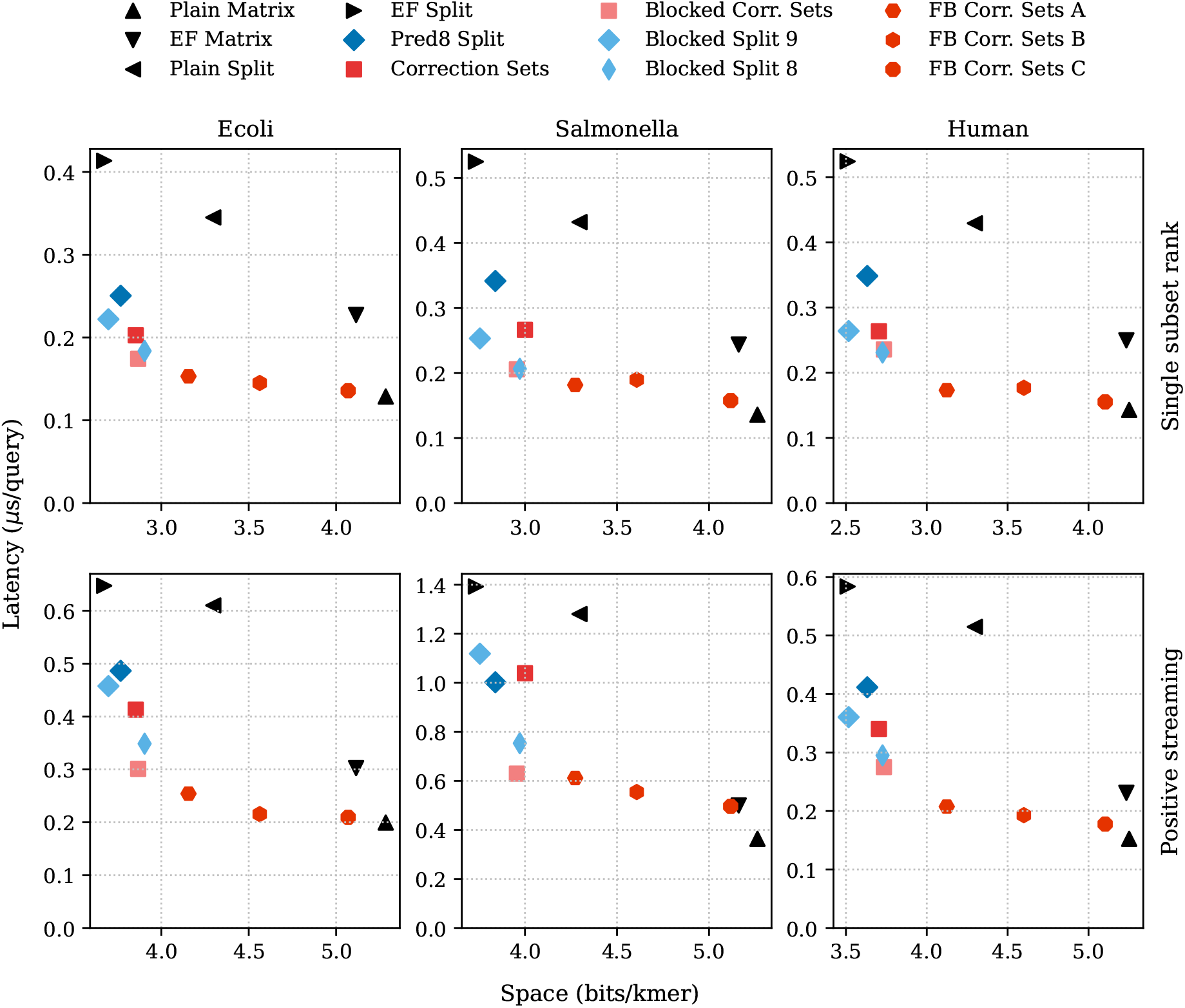
Latency and space usage for subset rank data structures on E.coli, Salmonella, and Human data sets. Top: latency (*µ*sec per query) over 10^6^ random single subset rank queries. Bottom: Latency (*µ*sec per query) over 10^5^ positive streaming *k*-mer lookup queries, *k* = 31. Space is 1 bitß higher per indexed *k*-mer in the bottom row of plots compared to the top plots because of extra space used to support streaming queries (see [3]).

Streaming k-mer lookup. Here we measure subset rank performance in the context of SBWT-based *k*-mer lookup, which uses subset rank queries intensively. *k* was set to 31 and *k*-mer lookup was performed in streaming mode (see [3]), where query *k*-mers come embedded in longer sequences (100,000 reads each of length 100bp in our experiments). We performed two separate experiments, one measuring the time for positive queries (see bottom of Fig. 5), where the reads are generated by sampling the underlying data, and negative in which the 100bp string is generated randomly and is unlikely to contain *k*-mer matches (see Appendix, Fig. S1).

There is an increase in latency for subset rank (top of Fig. 5) and positive *k*-mer lookup queries. This is expected because there are two subset rank queries performed per step during *k*-mer lookup (as explained in Section 2). However, the increase in the plots is less than a factor of two. This can be explained by cache effects: as a *k*-mer lookup query proceeds, the positions of rank queries become closer together and eventually start to hit a region of the subset rank data structure that falls within one cache line. With this in mind, we observe, however, that Matrix remains slightly faster than the other methods. We believe this can be explained as an effect of the computation involved in resolving a subset rank query *after* the relevant block is in cache being less in Matrix than in any of our blocked variants. In particular, to answer the second rank query, Matrix needs only to perform a popcount instruction on the same cache line it just accessed (in the first query), where as our blocked methods, must again scan through correction sets and perform a base-4 rank, which is more expensive than a popcount, even though the relevant data is also already in cache. This could be remedied by the implementation of a “subset rank pair” method, optimized to resolve queries at two nearby positions. We leave this as future work.

All-symbols subset rank. In this benchmark we generated 1 million random positions *i* and then queries for each of the four nucleotides, issuing four queries: subset-rank(*i*, A), subset-rank(*i*, C), subset-rank(*i*, G), and subset-rank(*i*, T). This benchmark is intended to simulate the kind of queries issued in the process of de Bruijn graph exploration, or in an approximate *k*-mer lookup, where more than one (even all) symbols are tested at a given step the lookup process. Results for the E.coli data set are shown in Fig. 6, right, along with single queries (left) and pairs of queries (center) for comparison. For these all-symbols queries, we now see some of the new Blocked subset rank structures overhaul Plain Matrix, which must access four distinctly different parts of memory (one for each of its bit vectors), whereas the blocked methods usually find the answers for all 4 queries in the same block.

**Figure 6.**
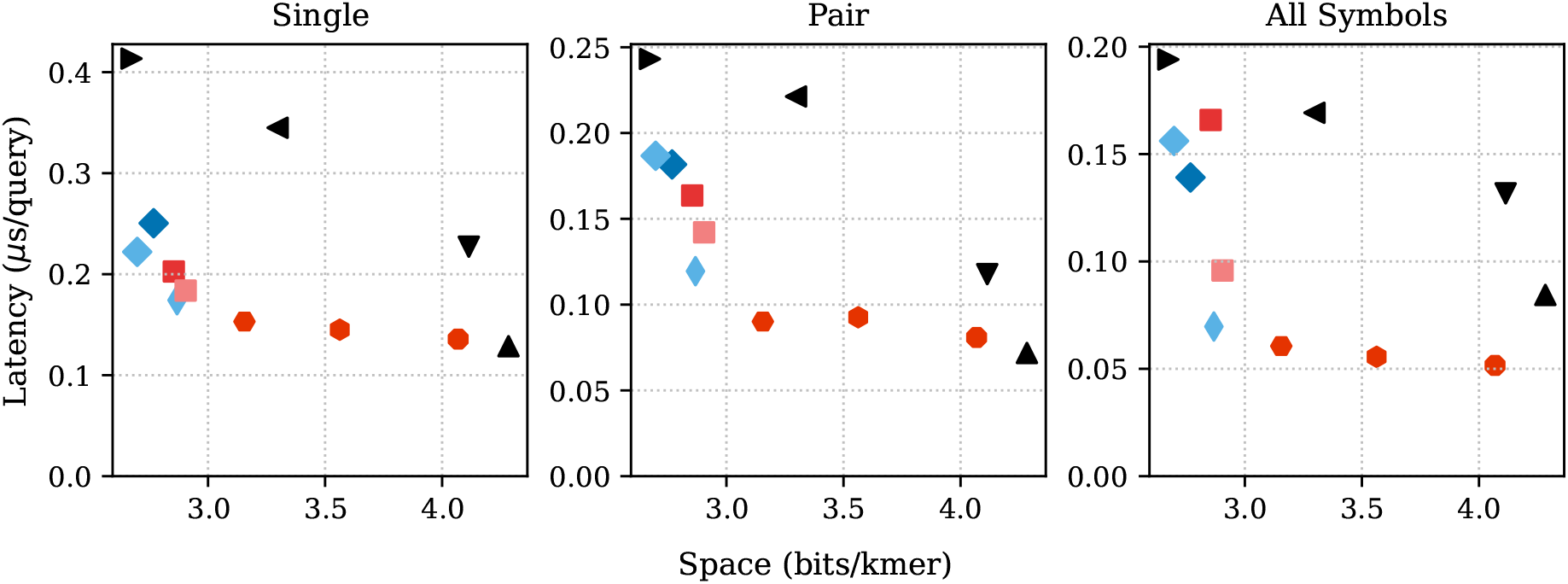
Results for different sets of subset rank queries on the E.coli dataset. Left: latency for 10^6^ single subset rank queries (replicating the top-left plot of Fig. 5). Center: latency for 10^6^ pairs of subset rank queries. Position *i* of the first query in the pair is random, the position of the second query is *i* + 5, with the same symbol. Right: Latency for 10^6^ sets of 4 queries at position *i* for all four symbols A, C, G, and T, with positions generated randomly.

Curiously, however, the difference is *not a factor of 4*, but at most a factor of 2. We suspect that over the long batch of queries the cache prefetcher is able to successfully detect and predict the constant “stride” of accesses made by Plain Matrix—despite being distant, they are spaced exactly *n* bits apart—and this reduces latency.

## 7 Concluding Remarks

We have described improved data structures for subset rank queries, which lead directly to faster small-space SBWT-based *k*-mer lookup. As a data structure, the SBWT enables a richer set of functionality than hashing-based *k*-mer lookup methods do, such as prefix search, and our methods translate into improvements for that functionality too. Our work in this paper has targeted the small-space end of the tradeoff curve. Our improvements have come from changes in design, to make data structure access more local, and in engineering of internal operations, such as base-4 rank. Any further improvements to base-4 rank data structures (e.g., [14, 13]) will lead directly to improved performance for our structures. We note that the reordering of bits we describe in Section 4.2 also appears in [13]. The possibility of improving on the speed of the Matrix representation, possibly by also increasing space usage, is an interesting avenue for future research. Another direction is to explore parallel processing of subset rank queries on our data structures in concert with batch processing [5]. The Correction Sets approach seems a particularly good candidate for multicore/GPU implementation because it lacks the data dependencies present in the Split structure.

## Acknowledgments

We thank Elena Biagi and Jarno Alanko, for frequent and wide-ranging discussions on the subject of the SBWT, and Ragnar Koerkamp for an interesting presentation and ensuing discussion at the Data Structures in Bioinformatics workshop in Venice, 18-19 February, 2026.

## A Other Data Structures

On our way to arriving at the data structures described in the paper, we had some less successful designs, which we include here, for completeness.

### A.1 A Split-like Concat structure without select

The main drawback of the Concat structure is the requirement of select support on the compressed encoding of the lengths of the subsets of *X*. In this section we propose new variants of Concat relying purely on rank supporting structures. The string *L* = *S*(*X*_1_)*S*(*X*_2_) … *S*(*X*_*n*_) is the concatenation of the characters of subsets *X*_*i*_ with support for base-4 rank. The empty sets are not included in the concatenation. A bit vector *B* of length *n* with *B*[*i*] = 0 if subset *X*_*i*_ is a singleton, and *B*[*i*] = 1 otherwise. Let 𝒞 be the set of lengths of subsets that are not singleton and let *m* be the size of 𝒞. A subset-rank(*i, c*) query is answered by *rank*_*c*_(*L, i*′), where *i*′ is the starting position of set *X*_*i*_ in *L*. If *j* = *rank*_1_(*B, i*) is the number of non-singletons before set *X*_*i*_ and *l* the sum of lengths in set 𝒞 up to and including position *j*, then *i*′ = *i*− *j* + *l*.

We index *L* with our base-4 rank structure, and *B* with a Pred8 structure. Next, we suggest two different ways to index set C to support prefix sum queries.

1. The four possible values {0, 2, 3, 4} in 𝒞 are concatenated and indexed using our base-4 rank data structure. Then *l* = 2 · rank_2_(𝒞, *j* + 1) + 3 · rank_3_(𝒞, *j* + 1) + 4 · rank_4_(𝒞, *j* + 1).
2. We exploit the property of SBWT that sum of lengths of must equal one less than the size of 𝒞. A vector *B*′ of length *m* indicates with bit *i* set to 1 that set *i* in 𝒞 is not of length 0. The lengths in 𝒞 that are not 0 are then concatenated to a binary string *L*′ using 2 bits per length. Length 2 is encoded as 00, length 3 encoded as 01, and length 4 as 11. The prefix sum of the lengths in 𝒞 up to position *j* is given by a rank query on vector *B*′, *b* = rank_1_(*B*′, *j*), and a rank query on bit string *L*′, *l* = rank_1_(*L*′, 2*b*) + 2*b*.

The space requirements for these structures are 2.25*n* bits for the concatenation of subsets *L* with supporting rank structures, *n/*8 + 8*m* bits for *B* and 2*m* for the encoding of the lengths of the non-singleton sets.

### A.2 A smaller version of the Split structure

Let *M*^−^, *M* ^+^ and *B* be as in Split from Section 3.2. We index *M*^−^ with our base-4 rank structure and *B* with a Pred8 structure. Let *B*′ be a bit vector of length *m*, where *m* is the number of columns in *M* ^+^. We set *B*′[*i*] = 1 if and only if column *i* of *M* ^+^ is not all zeros (i.e. *B*′ marks columns that do not correspond to empty sets). Let *M* ^++^ be the columns from *M* ^+^ that are marked with 1 in *B*′. We index *M* ^++^ as in the plain matrix representation. Splitting *M* ^+^ into *B*′ and *M* ^++^ instead of encoding *M* ^+^ using 4 bits per col reduces the space used for representing non-singleton sets by at least *m/*4 bits. *B*′ uses 1 bit per column and *M* ^++^ uses 4 bits for the remaining at most *m/*2 columns, in total *m* + 4*m/*2 = 3*m*.

Similar to split, a subset-rank(*i, c*) query on this new structure is answered by first issuing a rank query on *B*, then another rank query on *B*′ to determine how may of the *M* ^+^ were not all-zero columns. Finally, the rank structures of *M*^−^ and *M* ^++^ are used to count the number of characters in the corresponding prefixes of both matrices, and the sum of the two counts is returned. That is, if *r* = rank_1_(*B, i*) and *r*′ = rank_1_(*B*′, *r*), then the answer to the query is rank_1_(*M* ^++^[*c*], *r*′) + rank_*c*_(*S, i* − *r*).

## B Results for Negative Streaming *k*-mer Lookup Queries

We performed two separate *k*-mer lookup experiments, one measuring the per query latency for positive queries (see main text, bottom of Fig. 5), where the reads are generated by sampling the underlying data, and negative in which the 100bp string is generated randomly and is unlikely to contain k-mer matches, the results of which are shown in Fig. S1, below.

**Figure S1:**
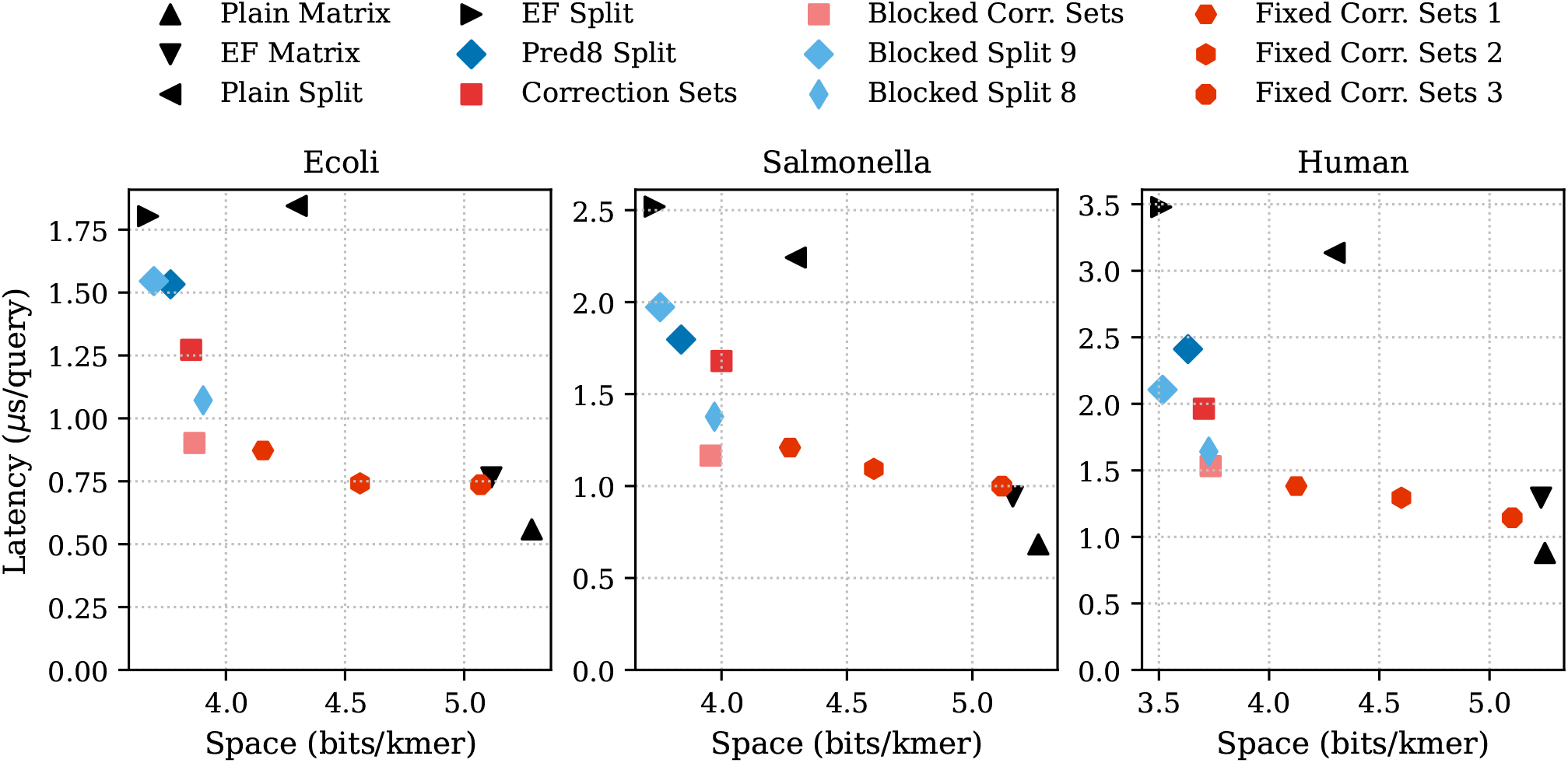
Results for negative streaming *k*-mer lookup queries on the E.coli, Salmonella, and Human datasets.

## Footnotes

1 Curiously, this arrangement resembles precisely the layout of bits in a balanced wavelet tree of the base-4 sequence.

2 The same is true for Concat, though the structures involved are different.

3 https://zenodo.org/records/6577997

4 The first 10,000 genomes from http://ftp.ebi.ac.uk/pub/databases/ENA2018-bacteria-661k/

5 The first 60 files from https://github.com/human-pangenomics/HPP_Year1_Assemblies

